# Tight hydrophobic core and flexible helices yield MscL with a high tension gating threshold and a membrane area mechanical strain buffer

**DOI:** 10.1101/2023.02.25.530059

**Authors:** Arjun Sharma, Andriy Anishkin, Sergei Sukharev, Juan M. Vanegas

**Author notes:** Correspondence: Juan M. Vanegas.

## Abstract

The mechanosensitive (MS) channel of large conductance, MscL, is the high-tension threshold osmolyte release valve that limits turgor pressure in bacterial cells in the event of drastic hypoosmotic shock. Despite MscL from *M. tuberculosis* (TbMscL) being the first structurally characterized MS channel, its protective mechanism of activation at nearly-lytic tensions has not been fully understood. Here, we describe atomistic simulations of expansion and opening of wild-type (WT) TbMscL in comparison with five of its gain-of-function (GOF) mutants. We show that under far-field membrane tension applied to the edge of the periodic simulation cell, WT TbMscL expands into a funnel-like structure with trans-membrane helices bent by nearly 70 degrees, but does not break its ‘hydrophobic seal’ within extended 20 μs simulations. GOF mutants carrying hydrophilic substitutions in the hydrophobic gate of increasing severity (A20N, V21A, V21N, V21T and V21D) also quickly transition into funnel-shaped conformations but subsequently fully open within 1-8 us. This shows that solvation of the de-wetted (vapor-locked) constriction is the rate-limiting step in the gating of TbMscL preceded by area-buffering silent expansion. Pre-solvated gates in severe V21N and V21D mutants eliminate this barrier. We predict that the asymmetric shape-change of the periplasmic side of the channel during the silent expansion provides strain-buffering to the outer leaflet thus re-distributing the tension to the inner leaflet, where the gate resides.

## 1 INTRODUCTION

Pressure or tension is a form of mechanical force that is central to many fundamental biological processes. The membrane-embedded mechanosensitive (MS) ion channels are responsible for converting mechanical forces into electrochemical signals that activate downstream cascades necessary to perform particular functions. MS channels are ubiquitous and play essential roles including elaborate sensory functions (Arnadóttir and Chalfie, 2010; Kefauver et al., 2020; Cox et al., 2018). In bacteria, MS channels from the MscS (MS channel of small conductance) and MscL (MS channel of large conductance) families fulfill the ancient housekeeping function of maintaining osmotic forces acting on the cell envelope within safe limits (Booth and Blount, 2012). In the event of a drastic osmotic down-shock, MscS and MscL channels open sequentially to release osmolytes and curb intracellular pressure. MscL is the high-threshold non-selective channel that opens at near-lytic tension as a last measure to rescue the cell from membrane rupture (Haswell et al., 2011).

MscL was the first identified and cloned MS channel directly activated by tension in the surrounding lipid bilayer. Its isolation from *E. coli* was reported in 1994 (Sukharev et al., 1994) and since then, a vast amount of functional data has been collected specifically on this channel species (Häse et al., 1995; Blount et al., 1996; Ou et al., 1998; Yoshimura et al., 2001; Ajouz et al., 2000; Bartlett et al., 2006). Attempts to crystallize several MscL homologs from different bacteria resulted in the structure from *M. tuberculosis* (TbMscL), which was solved in its closed state (Chang et al., 1998; Steinbacher et al., 2007). TbMscL shares ~30% identity with E. *coli* MscL (EcMscL) and has similar domain organization. MscL is made from a pentameric assembly with two transmembrane (TM) helices and both N- and C-termini on the cytoplasmic side. TM1 helices from each subunit line the inner pore and form a tight hydrophobic constriction on the cytoplasmic side. TM1 and TM2 helices are swapped between the neighboring subunits resulting in a stable design in which the periplasmic loops interlink all subunits, thus keeping the complex together under extreme tensions (Iscla and Blount, 2012; Chang et al., 1998; Steinbacher et al., 2007).

EcMscL is a much more convenient experimental system because it opens at less extreme tensions than TbMscL (Zhong and Blount, 2013). Yet, the crystal structure of TbMscL has been widely used as a common template for homology modeling (Sukharev et al., 2001a), prediction of gating transitions (Perozo et al., 2002), informed mutagenesis (Ou et al., 1998; Anishkin et al., 2005; Chiang et al., 2005), and extensive molecular dynamics (MD) simulations (Gullingsrud and Schulten, 2003; Jeon and Voth, 2008; Deplazes et al., 2012; Martinac et al., 2017). Early attempts at modeling suggested that the opening transition proceeds through tilting followed by radial outward motion of the TM1-TM2 helical pairs rather than forming a parallel ‘barrel-stave’ arrangement of the helices (Sukharev et al., 2001b; Betanzos et al., 2002). Thus far, none of the complete MscL homologs has been captured in a fully open state in any of the structures, leaving experimental estimations of spatial and energetic parameters to hypothesize the opening transition mechanism, which constrain the models, and delegating the atomistic features to detailed MD simulations.

Analysis of patch-clamp traces gave sufficient information about the number of functional states, energetic parameters and the character of helical movements in MscL. In giant bacterial spheroplasts and reconstituted liposomes, EcMscL behaves like a two-state (one-barrier) system with a number of short-lived subconductances surrounding the open state (Chiang et al., 2004). It is now a consensus that EcMscL’s tension midpoint for activation is near 12 mN/m (Sukharev et al., 1999; Moe and Blount, 2005; Nomura et al., 2012). Thermodynamic (Boltzmann) analysis of slopes of open probability on tension predicted a ~20 nm^2^ in-plane expansion associated with opening (Chiang et al., 2004). EPR (electron paramagnetic resonance) accessibility measurements (Perozo et al., 2002) and FRET (Förster resonance energy transfer) studies (Corry et al., 2010) gave consistent results and confirmed this spatial scale. Measurements of opening and closing rates as a function of tension suggested the position of the transition state on the protein expansion scale, a widely used reaction coordinate, at ~0.65-0.7 between the fully closed (0) and fully (1) open states. This indicated that the rate-limiting barrier is skewed toward the open state on the expansion scale that makes the well for the closed state wider and ‘softer’ (Sukharev et al., 1999; Chiang et al., 2004). The shape of the energy profile thus implied that the channel undergoes a substantial ‘silent’ expansion (13 nm^2^) before it transitions to the open state. With the aid of ultrafast patch-clamp, it has been shown that the actual transition between the closed and fully open states in MscL is shorter than 3 μs although the waiting time under constant super-threshold tension can be milliseconds and seconds until the opening barrier is overcome (Shapovalov and Lester, 2004).

There have been a number of mutagenesis studies targeting different domains in MscL (Yang et al., 2012; Yoshimura et al., 1999), but even an early unbiased random mutagenesis study gave unequivocal indication that the hydrophobicity of the MscL pore constriction is a prerequisite for the stability of the firmly closed and leak-proof resting state of the channel (Ou et al., 1998). Subsequent studies confirmed that essentially the same polar substitutions in the pore constriction found in the unbiased genetic screens de-synchronize the sequence of gating transitions and thus make accessible a multitude of subconductive states that become populated at low membrane tensions (Anishkin et al., 2005). This behavior was attributed to abnormal hydration and pre-expansion of the channel gate that is normally completely desolvated (vapor-locked) in the WT (Anishkin et al., 2010).

From a biological point of view, the cellular osmotic permeability response is a costly operation. Light scattering stopped-flow experiments performed on live bacteria estimated that abrupt dilution of the medium may lead to the release of ~15 % of all non-aqueous components from the cell, including hardly replenishable metabolites (Çetiner et al., 2017). This suggests that the sequential action of MscS and MscL should be somehow coordinated to minimize losses. With that regard, an interesting hypothesis was published by Joos and coworkers (Boucher et al., 2009) stating that ‘soft’ mechanosensitive channels present in sufficient density in the cytoplasmic membrane of *E. coli* may delay or avert cell lysis even without opening, just by acting as a membrane area buffer.

In the present work we perform multi-microsecond atomistic MD simulations (>45 μs combined) of TbMscL under membrane tension in two different regimes, (1) with far-field tension applied to the entire simulation cell and (2) using the locally distributed tension (LDT) protocol focusing forces on lipids immediately surrounding the protein (Rajeshwar T et al., 2021). To avoid ambiguities of homology models, we resorted to simulations of the genuine TbMscL crystal structure. Our data reveals that the opening pathway of WT TbMscL is interposed by a long-lived and highly expanded intermediate, which may act as a membrane area buffer. The stability of this funnel-shaped asymmetric intermediate, where the periplasmic side significantly expands, under extreme tension is defined by the completely desolvated hydrophobic gate and flexible helices. We also show that five gain-of-function mutants carrying polar substitutions in the gate briefly transition into similarly expanded non-conductive structures but fail to sustain these conformations for longer periods of time and fully open within microseconds. We discuss the possible implications of this opening pathway and the roles of gate desolvation and the outer rim expansion for MscL function.

## 2 METHODS

All simulations were conducted with the GROMACS molecular simulation package version 2021 (Abraham et al., 2015). Lipids were modeled with the GROMOS 43A1-S3 (Chiu et al., 2009) forcefield (FF), while the GROMOS 54A7 FF (Schmid et al., 2011) and SPC/E model (Berendsen et al., 1987) were used for protein residues and water respectively. The classical leapfrog integrator was used to calculate Newton’s equations of motion with a time step of 2 fs, and coordinates were recorded into the trajectory file every 5 ps. The LINCS algorithm was applied to constrain the lengths of all covalent bonds. A plain cutoff value of 1.6 nm was used to compute Lennard-Jones interactions. The long-range electrostatic calculations were calculated with the particle-mesh Ewald method using a real-space cutoff of 1.6 nm and Fourier grid spacing of 0.15 nm. The Nosé–Hoover thermostat was used to maintain a constant temperature of 37°C. The pressure was held constant at 1 atm with a semi-isotropically coupled Berendsen barostat for equilibration and Parrinello-Rahman barostat for production simulations in the absence of tension. The Berendsen barostat was used for constant high membrane tension simulations due to technical limitations in GROMACS.

The 1-palmitoyl-2-oleoyl-sn-glycero-3-phospho-ethanolamine (POPE) membrane bilayer composed of 50 lipids per leaflet was generated using the MEMGEN server (Knight and Hub, 2015). The membrane was equilibrated prior to embedding the crystal structure of wild-type (WT) TbMscL from *M. tuberculosis* (PDBID 2OAR, (Steinbacher et al., 2007)). The gain-of-function mutants V21T, V21N, V21A, V21D, and A20N were similarly embedded into the equilibrated POPE membrane. For the embedding process, high lateral pressure of 10000 bar was applied for 1 ns onto the POPE membrane using the method described by Javanainen et al. (Javanainen, 2014). The embedded TbMscL-POPE systems were solvated with a water layer thicker than > 1.5 nm between the protein and its periodic image in the z-dimension.

For all WT TbMscL and gain-of-function (GOF) mutants, the solvated systems were first energy minimized for 500-1000 steps using the steepest descent algorithm followed by a 300 ns simulation at constant temperature and pressure. The last frame of the *NPT* simulation was used as the starting point for a high tension run of 50 mN/m. This high tension accelerates the gating process, yet the membrane may become unstable and rupture after some time. Therefore, we closely monitor the integrity of the membrane/protein and stop the simulation once the system appears to becomes unstable and the area begins to increase uncontrollably. We then take a snapshot from a few nanoseconds before rupturing and use this snapshot to continue the simulation with a 5 mN/m lower tension. The same process is repeated as needed until the pore for each system has reached the expected fully open configuration (radius of 1.0-1.5 nm). For the GOF mutants, the final tension is reduced to 40-45 mN/m before reaching that state.

In the case of WT TbMscL, the channel pore did not expand significantly despite simulating for more than 20 μs at high membrane tension. Therefore, we applied an additional stimulus using the locally distributed tension (LDT) MD method described in our previous work (Rajeshwar T et al., 2021) to accelerate the full opening. The LDT MD method uses a biasing force concentrated on the lipids surrounding the protein channel (in the membrane plane) through a specially defined collective variable (CV), *ξ*. This CV is defined by a smooth hyperbolic tangent stepping function

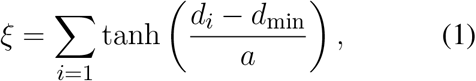

where *d_i_* is the lateral distance from the center of mass of each lipid to the center of mass of the protein channel, *d*_min_ = 1.1 nm is the minimum distance from the center of mass of the protein channel, and the constant *a* =1 determines how rapidly the hyperbolic function reaches unity. The system was biased with a linearly moving harmonic restraint or constant velocity pulling method where the biasing potential,

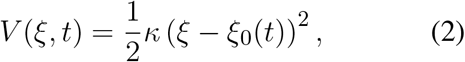

changes with time as the equilibrium position, *ξ*_0_(*t*), is gradually adjusted at a constant rate (*v*), *ξ*_0_(*t*) = *ξ*_0_(0) + *vt*. The WT TbMscL system was biased with LDT-MD over a period of 100 ns. A spring constant of *κ* = 100,000 kJ ·mol^-1^ · nm^2^ was used for the harmonic potential. The LDT MD steered simulations (Eq. 1) were performed using a PLUMED (Bonomi et al., 2009; Tribello et al., 2014) patched version (v. 2.7) of GROMACS 2021.

### 2.1 Data analysis

The pore radius at residue 21 was obtained from the subunit-averaged distance between the center of mass (COM) of each residue to the combined COM of all five residues in the plane of the membrane (*x – y*)

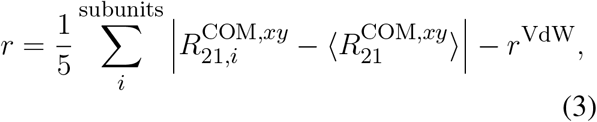

where *r*^VdW^ nm is the effective VdW radius for the residue (e.g., 0.29 nm for Val). The in-plane-projected protein area was calculated by creating a 2-dimensional Voronoi tessellation of all the protein and lipid atoms according to their *x – y* positions. The total protein area was computed by taking the sum of the individual areas of all protein atoms. Voronoi tessellations were calculated using the `gmx_LS voronoi` utility, which is part of the GROMACS-LS package (Vanegas et al., (accessed July 14, 2020) and is based on the Voro++ library (Rycroft, 2009).

The number of water molecules in the vicinity of residue 21 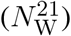 was obtained by finding any water that has an atom within 0.5 nm of any atom of residue 21 from any of the five subunits of the WT or GOF channels. The subunit-averaged transmembrane helix 1 bend angle, 〈*θ*_1,M_〉, was measured based on the vectors formed by the C*_α_* atoms of residues N13, A26, and I46, located at the beginning, middle, and end of the helix respectively. The bend angle is defined such that a straight helix has a value of 0.

Graphical representations of the MscL systems were created using UCSF Chimera and ChimeraX (Pettersen et al., 2004; Goddard et al., 2017). As there is some variability in degrees of tilting, bending and expansion among the five subunits of the TbMscL homo-oligomer, we performed an ‘instantaneous symmetrization’ of the WT TbMscL structures shown in Fig.8. This process helps to reveal the dominating trend and is for illustration purposes only. A symmetric average was first calculated using a custom-written Tcl script in VMD (Humphrey et al., 1996) from the coordinates of all five subunits at a given time point. After that, the protein atoms were energy minimized for 5,000 steps using conjugate gradient method to resolve all the structural conflicts, followed by a quick 1 ps simulation to ensure the absence of the clashes, and then symmetrized and energy minimized again.

## 3 RESULTS

### 3.1 Disrupting the hydrophobic lock in TbMscL

We investigate the pore expansion and wetting in TbMscL in multi-microsecond MD simulations by modulating the hydrophilicity of the pore at residues A20 and V21, which correspond to G22 and V23 in EcMscL. The WT channel as well as the GOF mutants A20N, V21A, V21N, V21T, and V21D were simulated in small POPE membrane patches (100 lipids total) and equilibrated for 300 ns in the absence of tension following energy minimization (see Methods). As tension-induced gating may take many microseconds depending on the hydrophobicity of the pore, we developed a variable tension protocol that facilitates gating while maintaining integrity of the membrane patch (i.e., preventing the bilayer from rupturing). For each system, we begin by applying a high system tension of 50 mN/m while carefully monitoring the channel pore radius at residue 21 as well as the overall system area (see Methods). We note that this high tension value, significantly higher than typical lytic tensions in the range of 10-15 mN/m, cannot be easily compared to experimentally measured values due to finite size effects that suppress long-range fluctuations in the small simulated membrane (see Marsh for a detailed discussion (Marsh, 1997)). Once the system becomes unstable and the area begins to increase uncontrollably, we stop the simulation and take a snapshot from an earlier time (few ns prior). We then use this time point to continue the simulation under tension lowered by 5 mN/m. The same process is repeated as needed until every channel pore reaches a radius of 1.0-1.5 nm (Fig. 1a). Our variable tension protocol produces a continuous high tension trajectory that results in all GOF mutants gating at tensions between 40-45 mN/m on a timescale of 0.5 to 4 μs.

**Figure 1.**
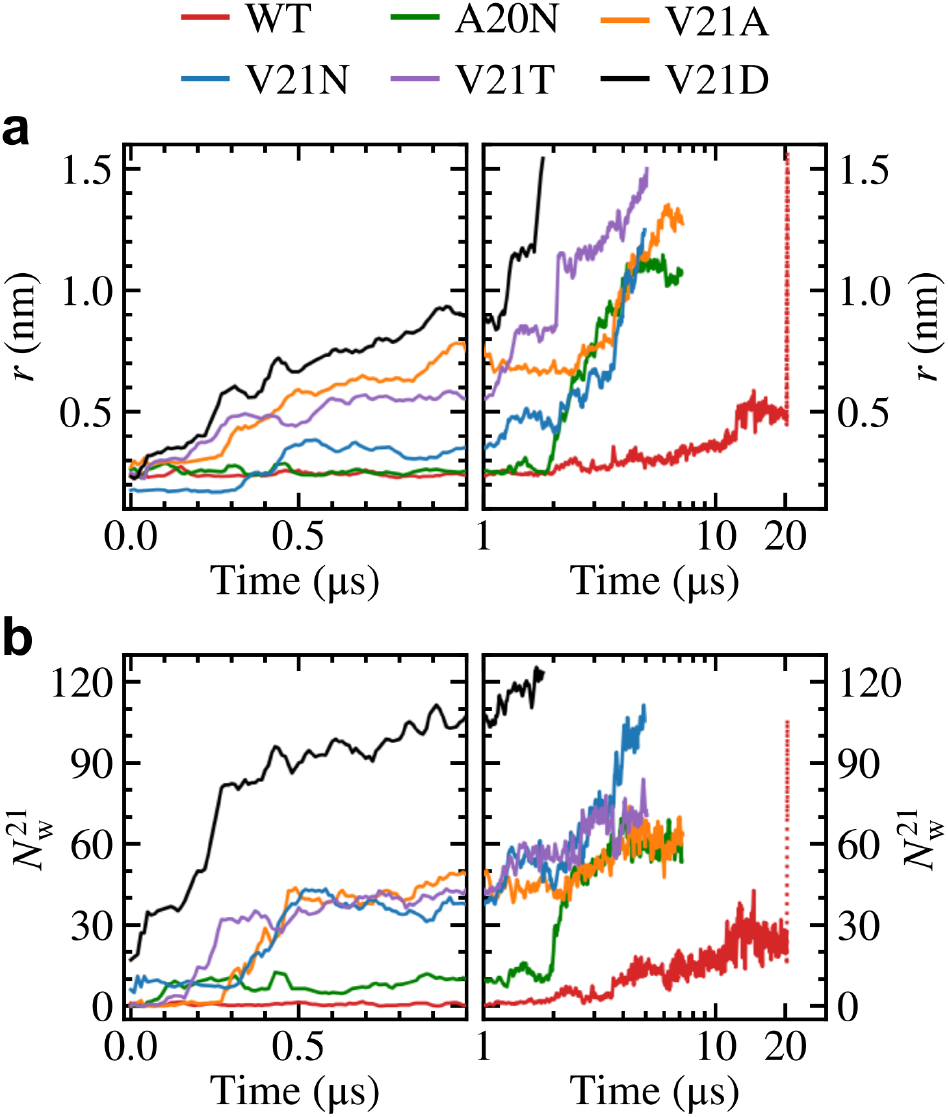
Gating WT TbMscL and GOF mutants under high membrane tension in multi-microsecond simulations. (a) Pore radius at residue 21 and (b) number of water molecules within 0.5 nm of residue 21. Left panels show time dependence at short time scales (*t* < 1 μs) while right panels show long time behavior (*t* > 1 μs) in a logarithmic scale. The red dotted line represents data for the WT system biased by LDT for 100 ns to accelerate pore expansion. The increased hydrophilicity of the GOF mutants facilitates wetting of the pore and allows faster expansion compared to the WT channel. The slow pore expansion in the WT results in intermittent wetting between 11 and 20 μs.

Focusing on the pore expansion of the GOF mutants first (Fig. 1a), we observe that V21D (black curve), which has the most drastic substitution due to its charged nature, shows the fastest pore expansion as its radius steadily increases the moment high membrane tension is applied. The pore radius for V21D reaches a value close to 1.0 nm after 1 μs and quickly climbs to 1.5 nm by 1.8 μs. A similar pattern is observed for the V23T mutant (Fig. 1a, purple curve), although the progression to the fully open state takes place at a slower rate. In contrast to these two mutants, the V21N channel (Fig. 1a, blue curve) remains tightly closed for the first few hundred ns and does not undergo significant expansion until after 2 μs. While the asparagine substitution at position 21 significantly increases the polarity in this region, hydrogen bonding between these residues in neighboring subunits makes the pore more tightly knit resulting in a smaller radius at *t* =0 and taking a longer time to gate compared to the V21D and V21T mutants. Introducing asparagine at the previous residue in the A20N mutant (Fig. 1a, green curve) keeps the pore radius virtually unchanged for the first ~2 μs under high tension, and then it begins to steadily increase until it settles near a value of 1.1 nm. The last GOF mutant tested is the V21A (Fig. 1a, yellow curve), which produces fast initial expansion of the pore within the first microsecond of the simulation similar to the V21D and V21T mutants despite alanine being a non-polar residue. After hovering for a couple of microseconds at a pore radius of ~0.8 nm, it continues to increase until it reaches a value of 1.3 nm after nearly 7 μs. Our simulation results are in excellent agreement with earlier experimental studies by Ou et al. (Ou et al., 1998) which observed that the equivalent mutations in *E. coli* (V23D, V23T, G22N, and V23A) resulted in “very severe” phenotypes making the cells unviable when the GOF mutants are expressed. Ou et al. also showed that the tension gating threshold is significantly reduced for E. *coli* V23A mutant, which opened before MscS in patch clamp experiments (Ou et al., 1998). The behavior of three GOF mutants of EcMscL has been studied experimentally (Anishkin et al., 2005). Hydrophilic substitutions for V23 (analog of V21 in TbMscL) de-synchronized the concerted opening and closing transitions by introducing multiple sub-conductive states. Analysis of transitions in these mutants indicated a pre-expanded closed state, apparently related to permanent and excessive hydration of the gate. Another series of experiments demonstrated that mutations in the periplasmic rim in EcMscL, on the other hand, impeded the channel opening and compromised its osmotic rescuing function (Yoshimura et al., 2004), consistent with the large conformational change on the periplasmic side.

In contrast to the GOF mutants, the pore radius for the WT TbMscL channel (Fig. 1a, red curve) does not change significantly for the first few microseconds and only moderately increases up to ~0.5 nm after 12 μs, where it hovers until the end of the simulation at 20 μs. As the WT channel may take many more microseconds to gate, requiring significantly more computing resources, we resorted to further bias the tensioned system at 20 μs using our previously developed locally distributed tension method (LDT-MD) over a period of 100 ns to expand the pore radius to ~ 1.5 nm (dotted red lines in Fig. 1).

We take a closer look at the hydration of the pore for the GOF mutants and WT by measuring the number of water molecules within 0.5 nm of residue 21 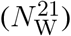 as shown in Fig. 1b. From this measurement, it is clear that the pore in the V21D mutant was well hydrated before application of tension 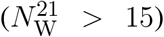 and the number of water molecules rapidly grows as the pore expands. The V21N mutant is the only other mutant which starts with a partially hydrated pore 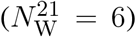 before tension is applied. Hydration of the pore in the V21N, V21T, and V21A mutants follows a similar pattern where the number of water molecules remains low for the first few hundred ns and then the number quickly grows to ~40 waters at *t* = 0.5 μs. In the case of the A20N mutant, there is an increase in the number of water molecules to ~10 after 100 ns, which is sustained up to 2 μs before growing to the final number of ~60 waters at the end of the simulation. Yet this initial partial hydration is not sufficient to completely wet the pore as it remains tightly closed during this 2 μs period.

Hydration of the WT channel follows a unique pattern where the ‘hydrophobic lock’ remains dry for the first few microseconds and the number of water molecules near V21 slowly grows between 2 and 20 μs. A close inspection of the TM region in the WT TbMscL shows that the narrow pore (~0.5 nm) becomes fully hydrated after 12 μs allowing water to connect the periplasmic and cytoplasmic sides as shown in Fig. 2. However, this wetting of the pore is unstable and it becomes dry again after a few nanoseconds (Fig. 2). This process is continuously repeated during the remainder of the 20 μs simulation as the pore flickers between wet and dry states. Despite the transient hydration with a few water strings, the WT TbMscL channel at this stage does not seem to be wide enough to allow a passage of hydrated ions through the hydrophobic gate and therefore is likely to be non-conductive until further expansion.

**Figure 2.**
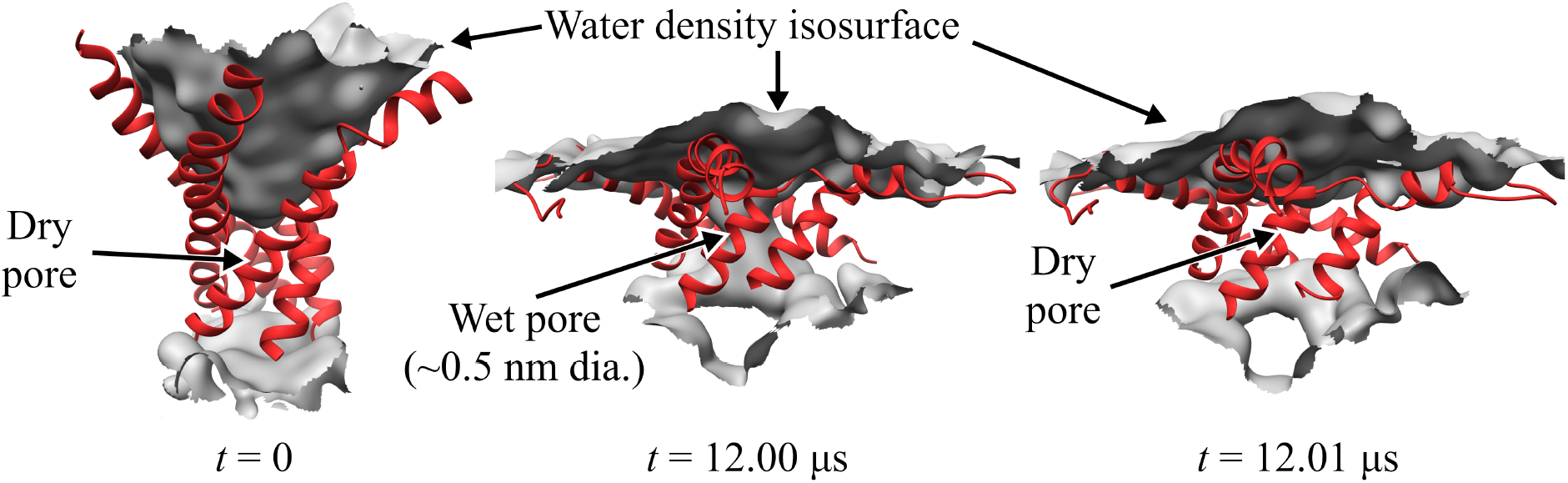
Close-up of WT TbMscL channel wetting/de-wetting during high membrane tension gating. TM1 helices shown as red ribbons and wetted regions shown through the isosurface of the water density (grey). All other protein elements and other molecules not shown for clarity. Before application of membrane tension, *t* = 0 μs, the hydrophobic residues of TM1 are tightly packed and the pore is dry (left panel). After applying high tension (50-45 mN/m) for 12 μs, the TM1 helices bend and the hydrophobic pore undergoes sufficient expansion to allow wetting and connect the periplasmic and cytoplasmic regions through a narrow water network (middle panel). The narrow wet pore is not fully stable and quickly re-closes after a few nanoseconds (right panel). The WT channel continues to ‘flicker’ between wet and dry states for multiple microseconds (see Fig. 1).

Visual examination of the initial (*t* < 0.5 μs) pore expansion of the GOF mutants provides further spatial insights into the process (Fig. 3). Beginning with A20N, we see that the introduced asparagine recruits numerous water molecules on the cytoplasmic side of the pore at *t* = 0 and this number grows quickly as tension is applied. However, the hydrophobic V21 keeps these cytoplasmic waters from wetting the pore for an extended period of time. In the case of the V21A mutant, the mildly hydrophobic alanine does not attract any water molecules, but its small size may facilitate the sliding of the TM1 helices past one another resulting in the relatively sudden gating of the pore between 0.25 and 0.5 μs. It is also possible that the small alanine sidechain cannot prevent the formation of a watery connection between the periplasmic and cytoplasmic aqueous compartments, which stabilizes the partially open state and eliminates the high barrier. As expected, the three polar/charged V21N/T/D mutants show a more progressive recruiting of water molecules and expansion of the pore as tension is applied. Beyond hydration of the pore, Fig. 3 shows that the WT and GOF mutants undergo significant bending of TM1 under tension. We further characterize this bending in the following section and its importance in relation to the pore radius and area expansion during membrane stretching.

**Figure 3.**
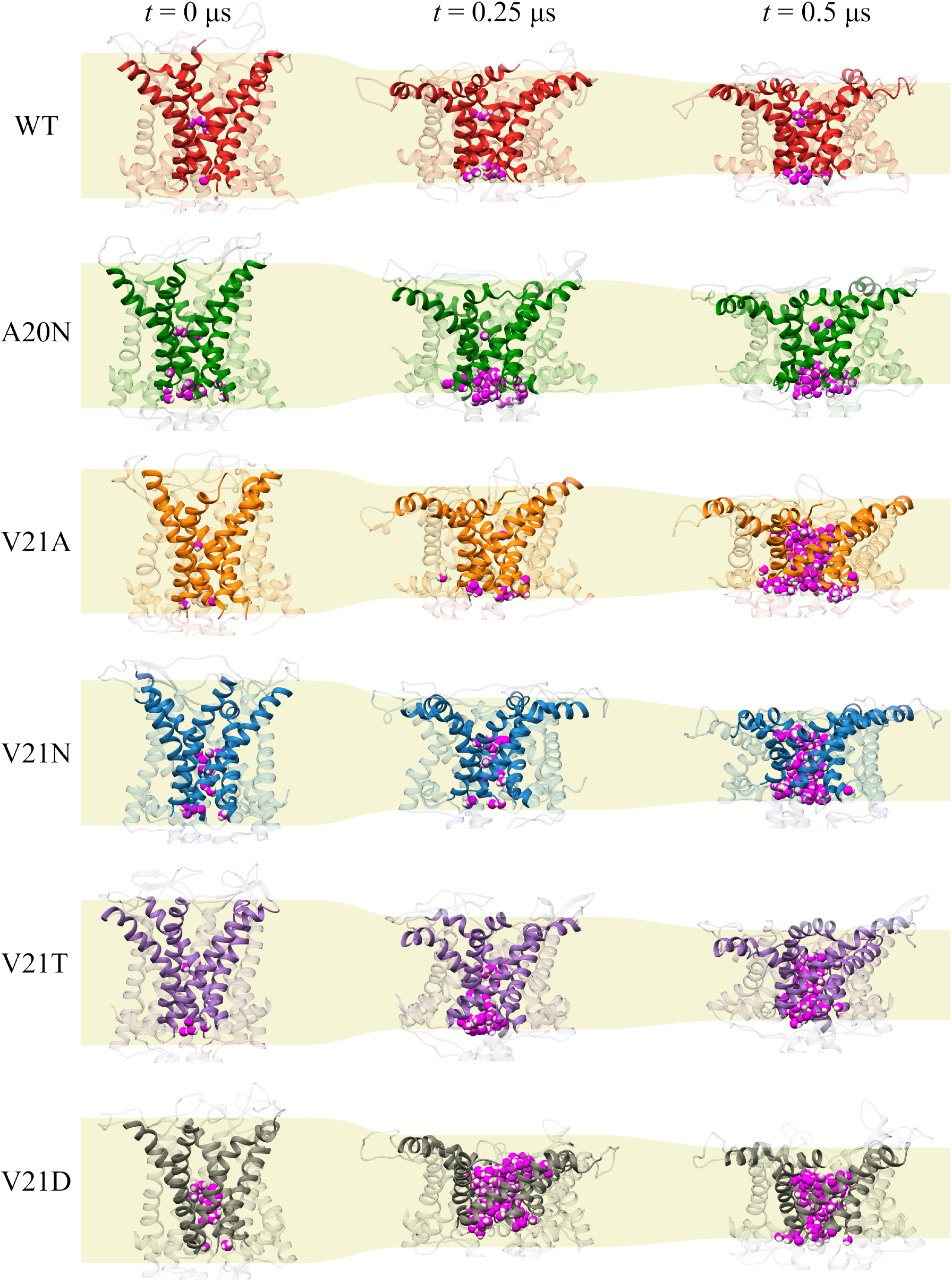
Hydration of of the inner pore and structural changes in WT TbMscL and GOF mutants during gating at high membrane tension. Ribbon representations of channels at rest (*t* = 0 μs, left) and two subsequent times, *t* = 0.25 μs (middle) and *t* = 0.5 μs (right). TM1 residues shown in solid colors while the TM2 and periplasmic loop residues are shown in transparent colors for clarity. Water molecules within 0.4 nm of any atoms of residues 20 and 21 are shown in magenta. The light yellow background roughly delineates the location of the lipid membrane as a guide to the reader. TM1 helices undergo significant bending under tension.

### 3.2 Bending of TM1 and silent expansion

We explore the functional role of TM1 flexibility on TbMscL gating by characterizing bending of the helix during application of high tension. The subunit-averaged TM1 bend angle, 〈*θ*_1,M_〉, is defined by the C*_α_* atoms of residues N13, A26, and I46 such that a straight helix has a value of 0. Rather than plotting the TM1 bend angle as a function of the simulation time, we present it as a function of the pore radius as shown in Fig. 4 in order to better characterize the structural features of the transition. In the WT channel (Fig. 4a) the pore radius remains unchanged as the TM1 bend angle reaches a value of 50 ° after which the pore moderately expands as the angle grows to its maximum value of 65 – 70 °. Once the WT pore expands beyond 0.6 – 0.7 nm, during application of the LDT bias in addition to tension (unfilled symbols in Fig. 4a), the TM1 bend angle gradually decreases to near its starting value (~ 35 °) as the radius grows to 1.5 nm. The A20N, V21A, and V21N mutants (Fig. 4b-d) show a similar pattern where their pore radius stays constant until the TM1 bend angle reaches threshold values of 45 – 50 °, which leads to an expansion regime where the radius increases while the angle remains roughly constant and later decreases. In the more severe hydrophilic V21T and V21D mutations (Fig. 4e,f) the pore expands soon after application of tension and therefore the initial bending of TM1 is accompanied by an increase in the radius, which is followed by a nearly constant or slowly decreasing bend angle as the radius approaches the maximum value of 1.5 nm.

**Figure 4.**
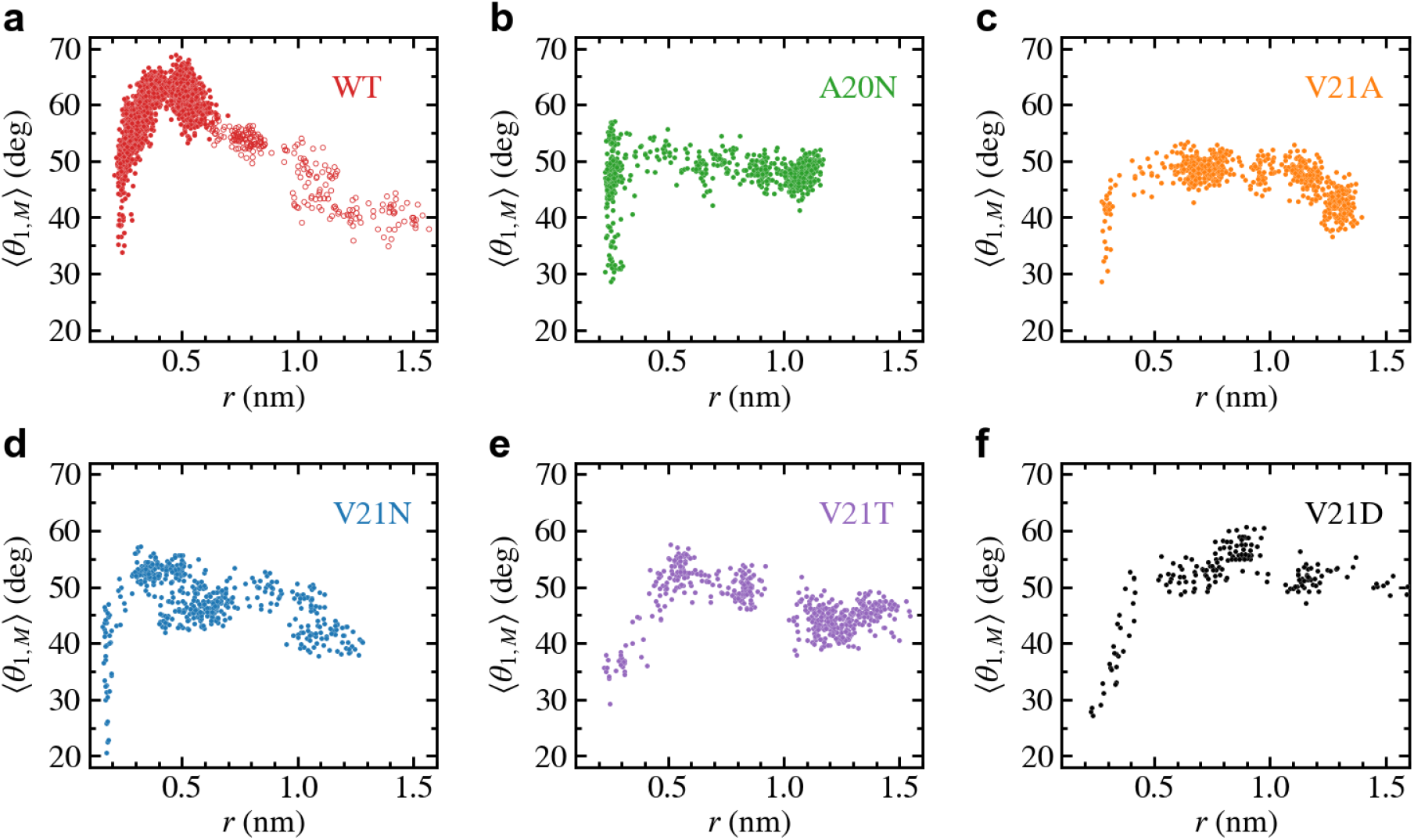
Bending of TM1 helix as a function of the pore radius for TbMscL channels under high tension. Data shown for WT (a) and GOF mutants A20N (b), V21A (c), V21N (d), V21T (e), and V21D (f). The subunit-averaged TM1 bend angle (〈*θ*_1,M_〉) is defined by the the C_α_ atoms of residues N13, A26, and I46 such that a straight helix has a value of 0. Unfilled circles in panel (a) show data for the WT system biased by LDT for 100 ns to accelerate pore expansion. Initial expansion of the pore is accompanied by a significant increase in the bend angle (> 50 °) for all GOF mutants and much more drastic for the WT (> 65 °)

As tension stretches the lipid membrane, the channel’s in-plane-projected protein area increases from a closed value of ~ 20 nm^2^ to ~ 40 – 45 nm^2^ in the fully open state as shown in Fig. 5. The fully open state was defined based on the pore radius increasing to 1.1-1.5 nm to satisfy the experimentally measured conductance of MscL. The rise in area of the WT (red curve) occurs more slowly compared to the V21 mutants as the hydrophobic lock keeps the inner pore closed yet the flexible protein elements allow for lateral expansion. The area of the A20N mutant, whose pore remains tightly closed during the first microsecond of the simulation, closely follows the values of the WT during this initial period of expansion. The relationship between pore radius and protein area for the WT and GOF mutants can be better appreciated when plotted versus one another as shown in Fig. 6. It is clear from this plot that the WT channel (Fig. 6a) undergoes a large ‘silent’ (i.e., non-conductive) expansion where the protein area increases by 15 nm^2^ or ~ 75 % with minimal change in the pore radius. This is in excellent agreement with previous experimental results that have shown that low subconductive states in EcMscL sustain large area expansions > 15 nm^2^ (Chiang et al., 2004; Anishkin et al., 2005). The silent expansion regime for the A20N mutant (Fig. 6b) is similar to that of the WT, but it is significantly reduced in the V21A and V21N mutants (Fig. 6c,d) where the pore radius begins to increase at a much lower area values. Area expansion in the V21T and V21D mutants (Fig. 6e,f) is always accompanied by an increase in the pore radius and therefore the silent regime is not observed.

**Figure 5.**
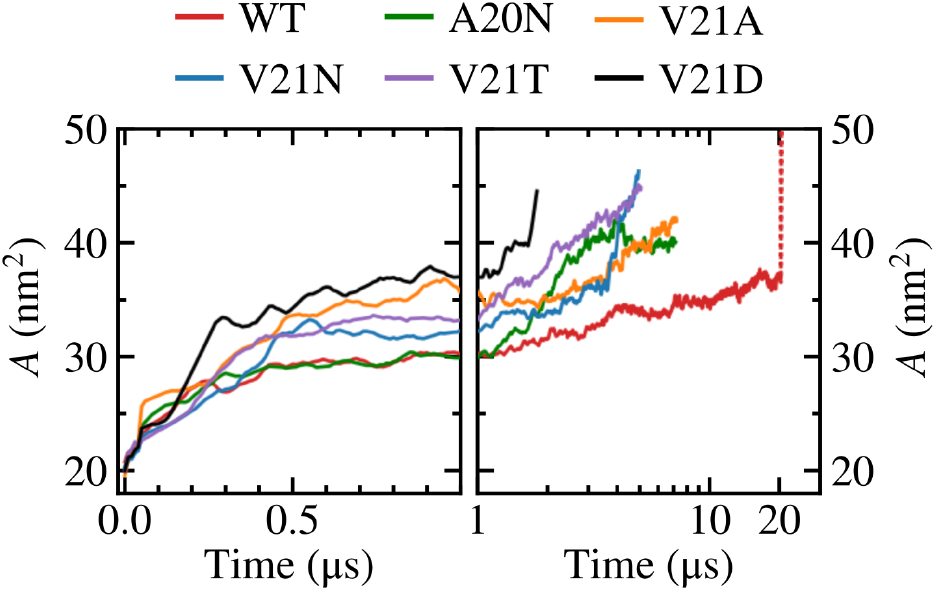
In-plane-projected protein area, *A*, of WT TbMscL and GOF mutants as a function of time. Left panels show time dependence at short time scales (*t* < 1 μs) while right panels show long time behavior (*t* > 1 μs) in a logarithmic scale. The red dotted line represents data for the WT system biased by LDT for 100 ns to accelerate pore expansion.

**Figure 6.**
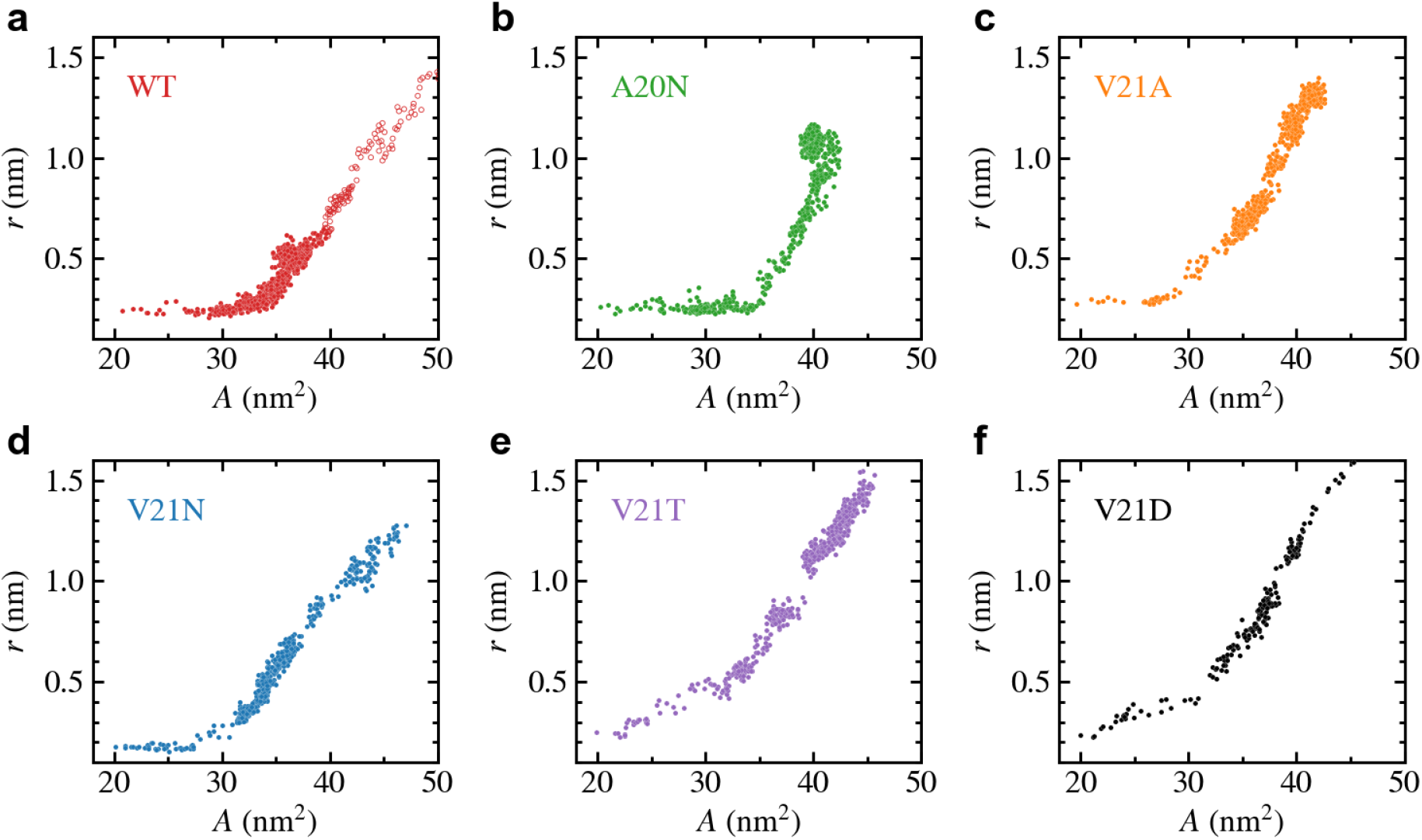
Change in pore-radius, *r*, as a function of the in-plane-projected protein area, *A*, for WT TbMscL (a) and GOF mutants A20N (b), V21A (c), V21N (d), V21T (e), and V21D (f). The subunit-averaged TM1 bend angle (〈*θ*_1,M_〉) is defined by the the C*_α_* atoms of residues N13, A26, and I46 such that a straight helix has a value of 0. Unfilled circles in panel a show data for the WT system biased by LDT for 100 ns to accelerate pore expansion. A silent expansion is observed for the WT and some mutants, while the more drastic mutations (V21T and V21D) show immediate increase in the pore radius as the protein area expands.

We further characterize the role of TM1 flexibility in the silent expansion by plotting the bend angle as a function of the protein area as shown in Fig. 7. The WT data (Fig. 7a) shows a unique pattern where the bend angle appears to linearly increase as the area expands up to a maximum value, which is followed by the reverse process where the area continues to expand while the TM1 helix straightens and the bend angle returns to near its starting value. At the bend angle peak, the WT protein adopts a funnel-like shape where the cytoplasmic side remains compact while the periplasmic side becomes widely spread. In the case of the GOF mutants (Fig. 7b-f), a similar trend is observed where the bend angle grows as the area increases and reaches a maximum value when the area is between 30-35 nm^2^. However, the decrease in the bend angle as the area increases past the peak point is much more gradual in the GOF mutants compared to the WT, yet the final bend is of similar size for all the systems.

**Figure 7.**
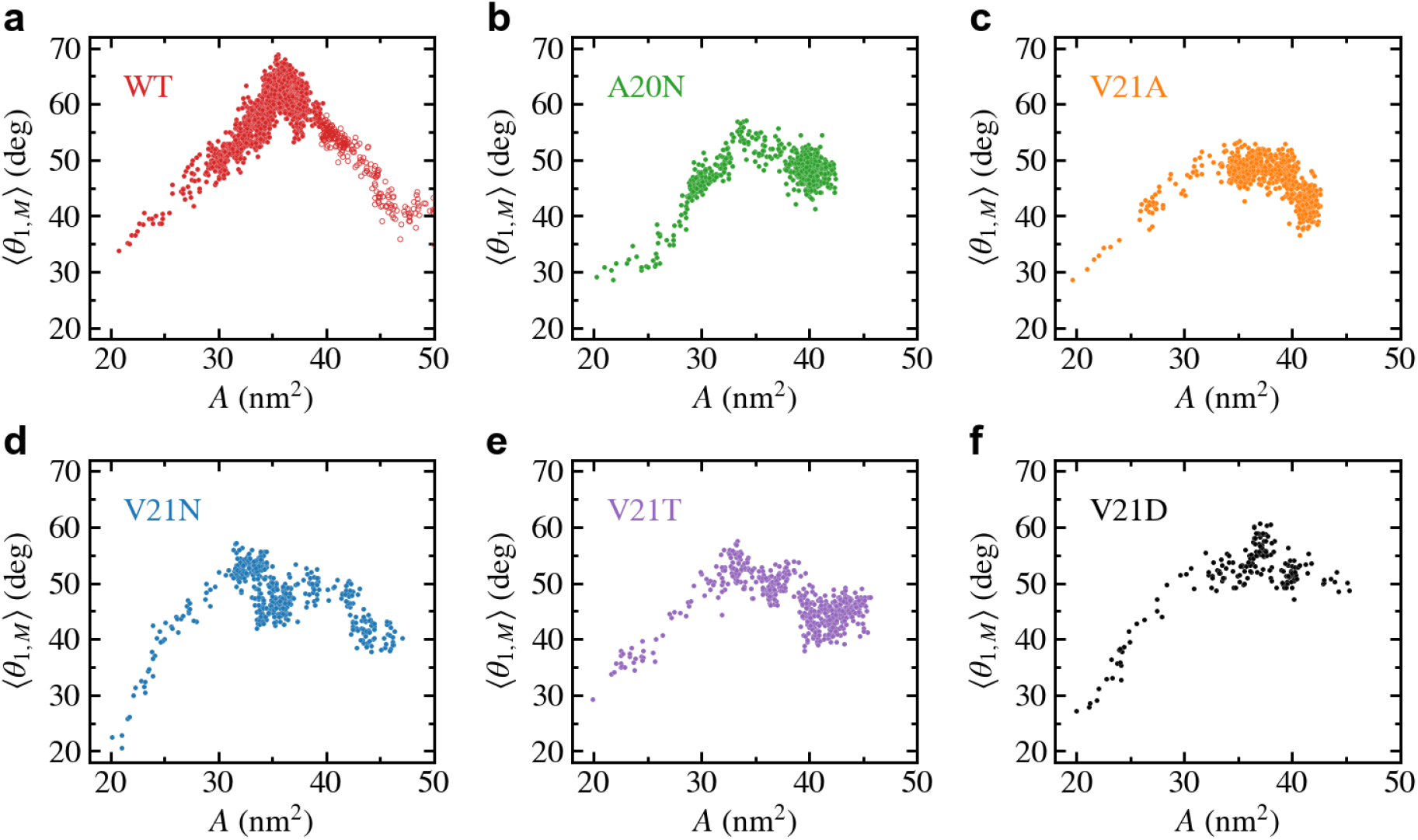
Bending of TM1 helix as a function of the in-plane-projected protein area, *A*, for TbMscL channels under high tension. Data shown for WT (a) and GOF mutants A20N (b), V21A (c), V21N (d), V21T (e), and V21D (f). The subunit-averaged TM1 bend angle (〈*θ*_1,M_〉) is defined by the the C*_α_* atoms of residues N13, A26, and I46 such that a straight helix has a value of 0. Unfilled circles in panel a show data for the WT system biased by LDT for 100 ns to accelerate pore expansion.

## 4 DISCUSSION

Our MD simulations of the TbMscL opening pathway performed under far-field tension are consistent with previous results and illustrate an asymmetric funnel-like expansion of the channel complex. This is illustrated in Fig. 8 where we show ‘symmetrized’ structures (see Methods) of the WT TbMscL before application of tension, in the tensed funnel-shaped silent intermediate, and finally in the fully open state after application of tension and LDT-MD bias. A wider expansion takes place at the periplasmic rim as the flexible TM1 bends while the inner pore near residues A20 and V21 remains tightly packed. In our simulations of the WT TbMscL, this expanded state is non-conductive as the hydrophobic gate remains completely dewetted for at least 20 μs of atomistic simulation under tensions of 50-45 mN/m. This result implies that the wetting step may introduce a substantial delay in opening experimentally.

**Figure 8.**
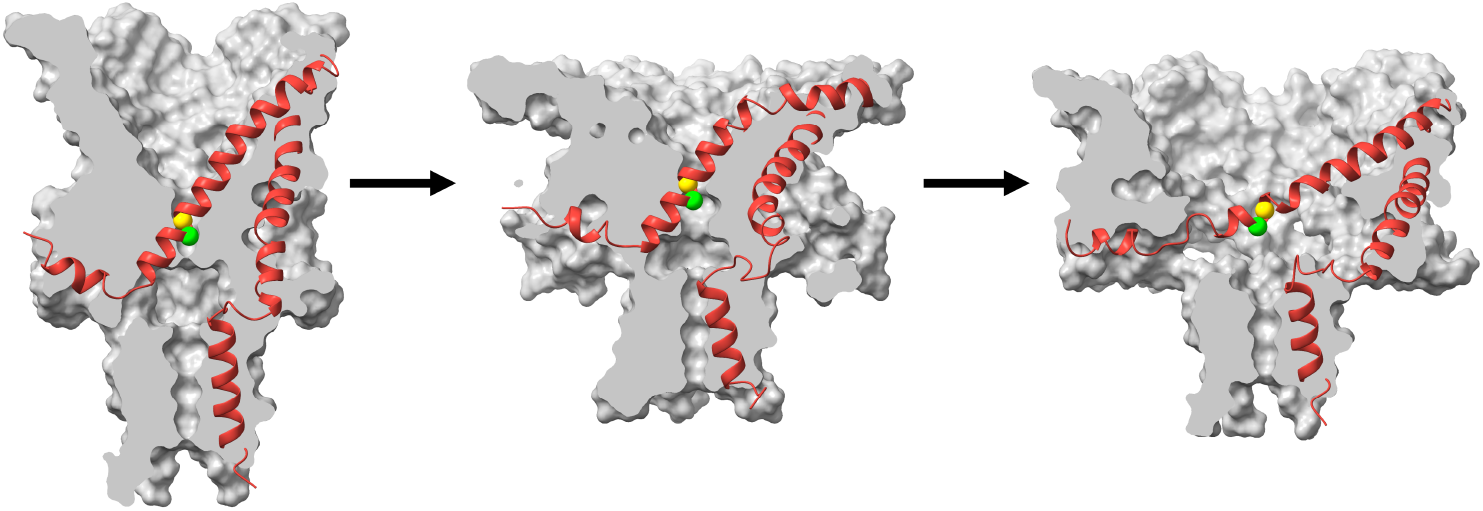
Cartoon illustration of the expansion and gating process in WT TbMscL during application of high tension. Surface cross section (grey) and ribbon representation (red) of a single subunit for symmetrized structures (see Methods) taken from the high tension simulation at *t* = 0 (left), *t* = 1 μs (middle), and *t* = 20 μs with additional LDT-MD bias (right). The green and yellow spheres show the C*_α_* atoms for residues A20 and V21 respectively. The channel first undergoes an asymmetric silent expansion where TM1 bends while the hydrophobic pore remains tightly packed. TM1 straightens and tilts as the pore expands to the fully open state.

As evidenced by the behavior of GOF mutants with hydrophilic substitutions in the gate, wetting of the hydrophobic constriction in WT TbMscL poses a rate-limiting barrier to the opening transition. The LDT-MD protocol helped us to push the WT channel system over this barrier and observe the full opening transition. Mutants with strongly hydrophilized pores opened earlier, which was accompanied by unimpeded wetting. In WT TbMscL, this barrier allows for the existence of a silently expanded state which acts as an elastic element and buffers the membrane area allowing time for other low-threshold channels, such as MscS, to act. As such, MscL opens when extreme tension is acting on the channel for a sufficient amount of time when other means to alleviate hypoosmotic stress are exhausted. GOF mutations lower this barrier and result in the almost immediate opening of the pore.

The strongly asymmetric character of expansion implies that this silent transition will buffer the strain specifically in the outer leaflet of the membrane. The expansion of the outer leaflet will lead to the re-distribution of tension to the inner leaflet and therefore pulling more efficiently on the gate. This mechanism may leverage (reinforce) the effect of tension on the cytoplasmically located channel gate. The data also predicts the interplay between the silent MscL expansion and gating of the small-conductance mechanosensitive channel MscS, which is also characterized by the cytoplasmic position of its gate.

## AUTHOR CONTRIBUTIONS

AS and JMV setup and carried out MD simulations. AS and JMV performed data analysis and created figures. AA provided conductance analysis scripts. JMV conceived the project and designed simulation studies with input from AS, AA, and SS. AS, AA, SS, and JMV contributed to writing of paper, reviewed, and approved it in its final form.

## ACKNOWLEDGMENTS

JMV and AS acknowledge the support of the National Science Foundation through Grant No. CHE-1944892. Computations were performed, in part, on the Vermont Advanced Computing Core supported in part by NSF Award No. OAC-1827314. This work used the Extreme Science and Engineering Discovery Environment (XSEDE), which is supported by National Science Foundation grant number ACI-1548562. XSEDE resources were provided at the San Diego Supercomputing Center (SDSC) Expanse system through allocation TG-BIO210110.

